# The plasma membrane bound transporter COPPER TRANSPORTER 2 facilitates palladium uptake in Arabidopsis

**DOI:** 10.64898/2026.09.17.752439

**Authors:** Jessica A. Dobson, Julia Jagielska, Frans JM. Maathuis, Neil C. Bruce, Elizabeth L. Rylott

## Abstract

Palladium (Pd), a technology critical platinum group metal, is increasingly lost to the environment through anthropogenic activity, yet the molecular basis of Pd uptake and tolerance in plants remains poorly understood. Here, we examine the role of the *Arabidopsis thaliana* (Arabidopsis) *COPT2* copper transporter in Pd transport and plant stress responses. Expression of *COPT2* in *Saccharomyces cerevisiae* conferred Pd sensitivity, indicating a capability for Pd transport. Arabidopsis *copt2* loss-of-function mutants accumulated less Pd in shoots and exhibited reduced reactive oxygen species (ROS) production, supporting a direct role for *COPT2* in Pd distribution and associated oxidative stress. Our subsequent transcriptome profiling revealed extensive changes under Pd exposure, including downregulation of *HMA2* and aquaporins, and upregulation of *HMA7*, glutathione transferases, and glutamine synthetase *GLN1;1*, consistent with the activation of metal detoxification and redox homeostasis pathways. Together, these findings identify *COPT2* as a key transport protein mediating the uptake of Pd in Arabidopsis and suggest that Pd induces similar detoxification mechanisms as zinc (Zn), copper (Cu) and cadmium (Cd), providing a foundation for engineering plants for precious metal recovery.

## Introduction

Palladium is used extensively in catalysis, particularly petroleum and chemical processing and electronics industries, and as jewellery, but the overwhelming use, >82%; 242 tons in 2022, is in catalytic converters for air pollution control in vehicles (Garside 2023). Reserves of Pd, and other platinum Group Metals (PGMs) are geographically concentrated, with 88% of current reserves in South Africa, and the rest in Russia, Zimbabwe, Canada and the United States. The technologies in which PGMs are used lack comparable substitutes, and PGM extraction and processing have high environmental impacts (Graedel *et al*. 2015). Thus, there is high demand, and supply chains are vulnerable. Sustainable circular systems for already-mined Pd offer a way to buffer and contribute to demand. While high levels of PGMs are recovered from catalytic exhaust systems at end-of-life (Saguru, Ndlovu and Moropeng 2018), during use, these metals are lost from the converters via exhaust fumes, with significant (up to 0.1 mg/kg) levels being deposited onto road and verge surfaces (Hooda *et al*., 2007). The process of mining and extraction of metals from primary ore is highly energy intensive and the volume of waste generated by mining is vast. As the ratio of ore to concentrate is typically 30-50:1, 96-98% of the ore becomes tailings (Glaister and Mudd 2010). In addition, there are slag wastes from smelters and other waste rock and water wastes. The resulting tailings dams and lagoons, often legacy or indefinite storage measures, can fail causing catastrophic environmental damage and loss of life (Yu *et al*., 2025).

Methods are urgently required to recover PGMs from dilute levels and remediate the large areas of land involved. Plants could be a potential solution in the form of phytomining. Phytomining harnesses the capabilities of specific plant species to concentrate metals from contaminated or geologically rich soils (Whitworth *et al*., 2022). Plant species have evolved to hyperaccumulate many metals including Zn, Ni and Cd, but to date no Pd hyperaccumulator species have been identified which is not surprising given its rarity and the abilityof Pd to readily form stable, inert complexes preventing plant uptake.

However, Pd uptake has been demonstrated in plants, with the metals forming nanoparticles *in planta* (Harumain *et al*., 2017; Kińska *et al*., 2019). Our research, and that of others (Harumain *et al*., 2017), has demonstrated downstream roles for Pd-rich plant biomass that add financial value to the phytoremediation and phytomining process beyond the use of plants to recover bulk metal. Harumain *et al*. (2017) used mustard (*Brassica alba*), and phytoremediation-relevant species, miscanthus (Miscanthus × giganteus) and willow (*Salix* sps.) to show the potential for Pd uptake from synthetic and mine-sourced tailings, and catalytic activity of Pd-rich pyrolysed biomass. Subsequent life cycle assessment on the phytomining approaches utilised, indicated that this technology has the potential to decrease the overall environmental impacts associated with extracting Pd using present day mining processes.

The *Arabidopsis thaliana* (Arabidopsis) COPT family of transport proteins comprises six members and plays a critical role in Cu homeostasis, with COPT2 (At3g46900) being involved in Cu transport under conditions of Cu deprivation (Mir, Pichtel and Hayat, 2021; Sancenón *et al*., 2003). Tiwari *et al*. (2017) confirmed the gene expression analyses by Taylor *et al*. (2014) that COPT2 has the capability to act as an Au transporter. A subsequent study demonstrated that Arabidopsis lines over-expressing COPT2 accumulated enhanced levels of copper (Cu) and Au in aerial tissues (Loskarn *et al*., 2024). Given the chemical similarities between Au and Pd, we hypothesised that Pd might also be a substrate for the COPT2 transporter in Arabidopsis.

## Materials and methods

### Yeast strains

Four *Saccharomyces cerevisiae* (yeast) strains were used in this work: BY4741 (*MAT* a *his3Δ1 leu2Δ0 met15Δ0 ura3Δ0*) and the isogenic mutant strain Y03629 (BY4741; *MATa*; *his3Δ1*; *leu2Δ0*; *met15Δ0*; *ura3Δ0*; YDR270w::kanMX4) were obtained from Euroscarf (Frankfurt, Germany). DTY165 (*MATα, ura3-52, his6, leu2-3-112, his3-D200, trp1-901, lys2-801, suc2-D* and isogenic mutant strain MPY17 (*MATα, ctr1::ura3::KanR, ctr3::TRP1, his3, lys2-801, CUP1*) were kindly donated by Dr Aaron Smith and Professor Dennis J. Thiele (Duke University, USA).

### Yeast spot dilution assay

Codon optimised *COPT2* was cloned into pYES2.1V5-His TOPO (Thermo Fisher Scientific) then transformed into yeast using the Frozen-EZ Yeast Transformation II Kit (Zymo Research). The presence of pYES2-COPT2y was confirmed using PCR. Transformed strains were grown overnight (30 °C) in medium containing 2 % (w / v) glucose, 6.7 g / L yeast N base without amino acids, and 1.9 g / L yeast synthetic drop-out medium supplements without uracil (SD-U; Sigma-Aldrich). Overnight cultures were centrifuged at3000 rpm for five minutes and washed twice with water, then resuspended and serially diluted with SD-U medium containing either 2 % galactose or 2 % glucose to an OD_600_ 1.0. The culture was spotted (3 μl) onto SD-U solid medium containing either 2 % (w / v) galactose or glucose, and additional metals (100 μM Cu, 500 μM Pd or 100 μM Cu + 500 μM Pd). Plates were incubated at 30 °C and images taken after five days using an Epson Perfection V370 Photo Flatbed scanner.

### Arabidopsis plant lines

The *copt2-1* (GABI-kat line ID: 168D01-013363) and *copt2-3* (GABI-kat line ID: 314H06.1) lines were identified from the German Plant Genomics Research Program (GABI-KAT; Kleinboelting et al., 2011). The *copt2-2* line (SALK_147451.31.40.x) was obtained from the Salk Institute Genomic Analysis Laboratory collections (O’Malley and Ecker, 2010). Seeds for *copt2-1, copt2-2* and *copt2-3* were obtained from the Eurasian Arabidopsis stock centre (N416069, N655195, N430138 respectively), along with the Columbia-0 (Col0), background ecotype.

Nucleic acids were extracted from two-week-old Arabidopsis rosette tissue using the EasyPure Plant RNA kit (Transgen Biotech co. LTD) with DNase Max® kit (Qiagen) to remove contaminating DNA; and the DNeasy PowerPlant Pro kit (Qiagen). The T-DNA insertion sites were determined following sequencing of PCR products using the primers described in Table S1. The qPCR was conducted as described in Tzafestas *et al*. (2017); *COPT2*-specifc, and constitutive *ACTIN2* primers used to normalise the data are listed in Table S1.

### Root response to Pd

Sterilised seeds were germinated and grown on solid agar (9g/L) plates containing half-strength Murashige and Skoog (1/2MS). Filter-sterilised Pd (as 50 mM K_2_PdCl_4_) or Cu (as 30 mM CuSO_4_) was added to concentrations shown in the figures, and pH adjusted to 5.7. Plants were grown with 16-hour day/8-hour night lighting at 100 μmoles m^-2^ sec^-1^ and temperatures of 24 °C/20 °C. Root length and number of branches of 12-day old plants were measured from images analysed using ImageJ. Root architecture was calculated as branches per mm (branches / length). Data were analysed using a two-way ANOVA, with a p-value of 0.02 due to deviations from assumptions in the data.

### Arabidopsis hydroponic system

Ten-day-old seedlings were grown as described above, then transferred to micropipette tip boxes containing 1/2MS, and six plants positioned using foam bungs within polystyrene rafts, as illustrated in Figure 5A. The plants were grown in a 12-hour day/night cabinet with 180 μmoles m^-2^ sec^-1^ and 21 °C/18 °C day/night photoperiod, with 1/2MS media refreshed every seven days. Five-week-old plants were dosed for 24h with 1 mM Pd, then root and shoot tissues were harvested and washed as follows:

### Preparation of samples for element analysis

Root samples were submerged in a metal desorption solution (2 mM CaSO_4_ and 10 mM ethylenediamine tetra-acetic acid) for ten minutes and shaken (180 rpm) to remove metals bound to the outside of the root tissue. All tissue samples were washed three times with 50 mL water and patted dry with tissue then dried at 90 °C for 72 hours, disrupted using a glass rod and washed, using 10mL of methanol, into glass 40mL vials and then sonicated for 20 minutes. The samples were then heated at 100 °C (with loose lids), for 8h, and the dry samples digested (tightly capped) overnight in 500 μL hot acid 1:3 molar nitric acid and hydrochloric acid at 110 °C. The cooled samples were diluted with water to a final volume of 10 mL in a 10 mL volumetric flask, then filtered using 1.6 μM pore Whatman Glass microfibre filters. The concentrations of Cu, Pd or Fe was determined by ICP-OES using an iCAP 7000 series (Thermo Fisher). Standard curves for these elements were generated using serial dilutions of Periodic table mix 1 and Periodic table mix 2 for ICP (Merck, UK). Palladium was measured at 340.269 nm, Cu at 213.598 nm, and Fe at 239.562 nm. Analysis was performed using a two-way ANOVA, using a p-value of 0.02.

### Analysis of ROS production in Arabidopsis root tips

Twelve-day-old plants, grown as described above, were transferred to six-well plates (Starlab) containing ½ MS medium with or without 20 μM Pd. After 24 h exposure, roots were washed once with 1× PBS (pH 7.4) and incubated in 10 μM 2′,7′-dichlorodihydrofluorescein diacetate (H_2_DCFDA; Sigma-Aldrich) dissolved in DMSO for 30 min. Samples were then washed four times with PBS before imaging. Oxidised dichlorofluorescein (DCF) fluorescence (λ_em = 522 nm) was visualised using a ZEISS LSM 710 epifluorescence microscope. Fluorescence intensity was quantified as corrected total cell fluorescence (CTCF), accounting for background signal (El-Sharkawey, 2016). To minimise root autofluorescence, images were captured using a lambda filter (9% laser exposure; 488 nm excitation; gain = 812). For FITC imaging (493–560 nm), 3% laser exposure and gain = 820 were used. A constant exposure time was maintained across all images.

### Transcriptome analysis

Five-week-old hydroponically grown Arabidopsis (Col-0) plants were exposed to 1 mM Pd for 24 h before harvest. Shoots and roots were separated, washed three times with sterile distilled water, and roots treated with metal desorption solution for 10 min before a final rinse. Five biological replicates (each comprising six pooled plants) were snap-frozen in liquid nitrogen. RNA was extracted, quantified, and quality-checked using a Qubit 4 Fluorometer. Poly(A)-enriched mRNA was reverse-transcribed and amplified for sequencing (Novogene, Cambridge UK) on an Illumina HiSeq platform. Reads were trimmed with CutAdapt v2.3 and quantified using Salmon v1.1.0 (TAIR10.51 reference; options: “validateMappings,” “seqBias,” “gcBias,” 100 bootstraps). Differential expression analysis was performed in DESeq2 v1.36.0, with gene annotation from the org.At.tair.db package v3.17.0.

## Results and Discussion

### Characterisation of S. cerevisiae expressing codon-optimised Arabidopsis COPT2 (COPT2y)

To characterise the involvement of COPT2 in Pd uptake, growth assays were performed with pYES2-COPT2y expressing strains on SD-U(A) media containing 100 μM Cu (Fig. 1A), 500 μM Pd (Fig 1B) or 100 μM Cu + 500 μM Pd (Fig. 1C). Sensitivity to Pd was assessed through comparison of growth on the galactose-containing medium to growth on a glucose-containing medium, as the negative control. Growth of both wild type strains (BY4741 and DTY165) expressing *COPT2* was severely inhibited by the presence of Cu, and to a lesser extent, by Pd. MPY17 has deletions in two, high-affinity copper transporter genes (*CTR1* and*CTR3*), which reduces Cu uptake. Thus, growth on Cu was enhanced in MYP17 transformed with *COPT2*, and, unaffected by the presence of Pd. Y03629 has a deletion in *Ccc2* which encodes a P1B-type ATPase that transports Cu into the Golgi apparatus, and thus a reduced ability to regulate intracellular Cu.

**Figure 1.**
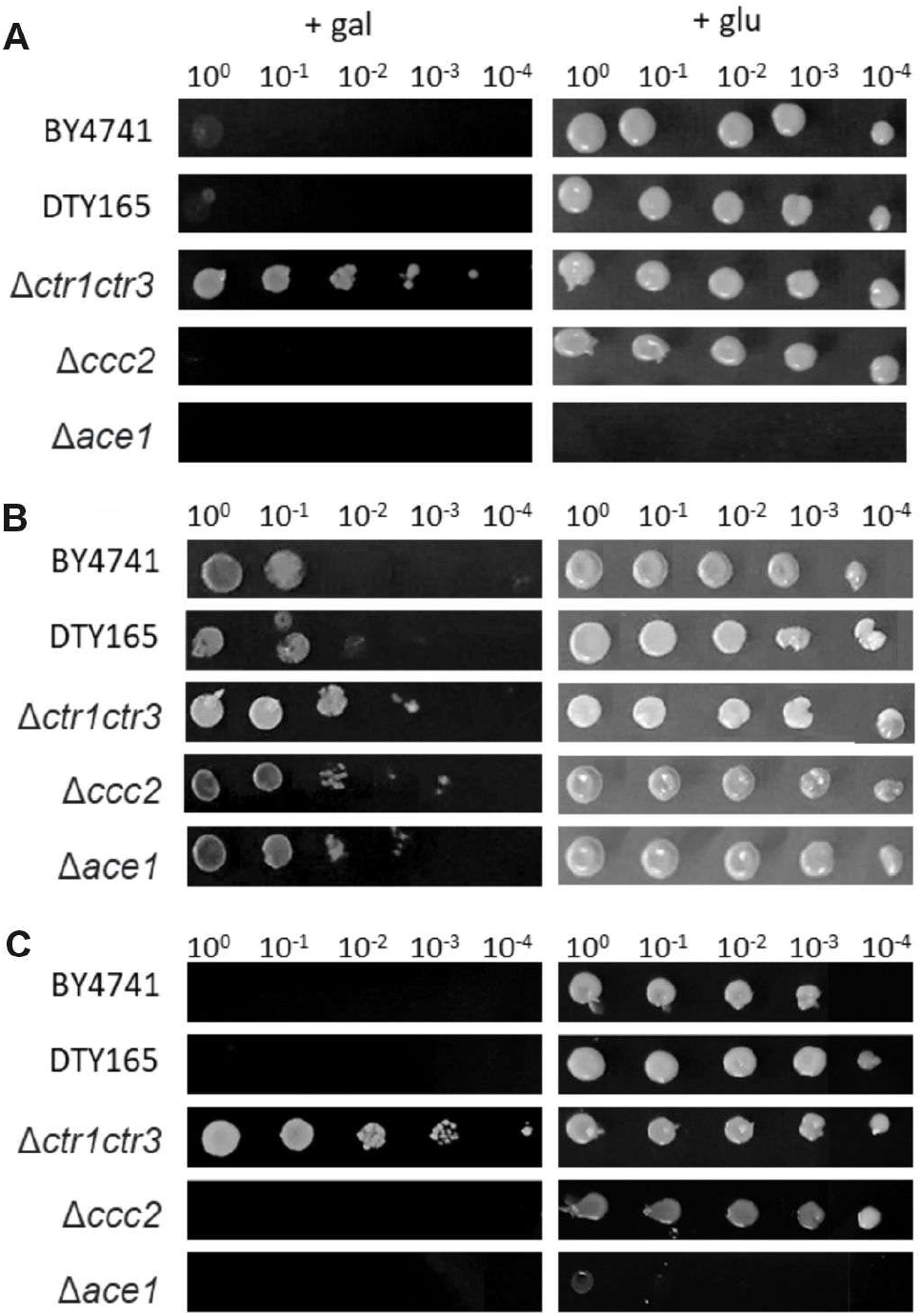
*Saccharomyces cerevisiae* strains expressing *COPT2y*. Spot dilution growth assay on SD-U(A) media containing Cu, Pd or both, using yeast strains containing pYES2-COPT2y. (A)100 μM Cu (B) 500 μM Pd, (C) 100 μM Cu + 500 μM Pd. The COPT2y sequence is under the control of the GAL1 promoter and therefore expression of COPT2y only occurs with galactose (gal) as the available carbon source. The SD-U(A) media, used for positive selection of yeast colonies containing pYES2-COPT2y, contained either gal (resulting in COPT2y expression) or glucose (glu) (no COPT2y expression). Cultures were grown to an OD_600_ of 1 and serially diluted by a factor of 10 (100, 10-1, 10-2, 10-3, 10 -4). Each row shown in the figure is a single strain and each ‘spot’ is a serial dilution of the one before, moving from left to right. Images are representative of the growth responses shown across five technical replicates.

### Characterisation of three Arabidopsis copt2 lines

Figure 2A illustrates the presence of the T-DNA insertion in all three lines. In *copt2-1* and *copt2-2*; 13 bp and 89 bp, respectively, upstream of the translation start site. In *copt2-3*, the T-DNA insertion is 90 bp downstream of the *COPT2* translation start site (Fig. 2B), with subsequent qPCR analysis (Fig. 2C) demonstrating that, in comparison to the wild type, *COPT2* transcript was undetectable in leaf tissue of all three *copt2* lines.

**Figure 2.**
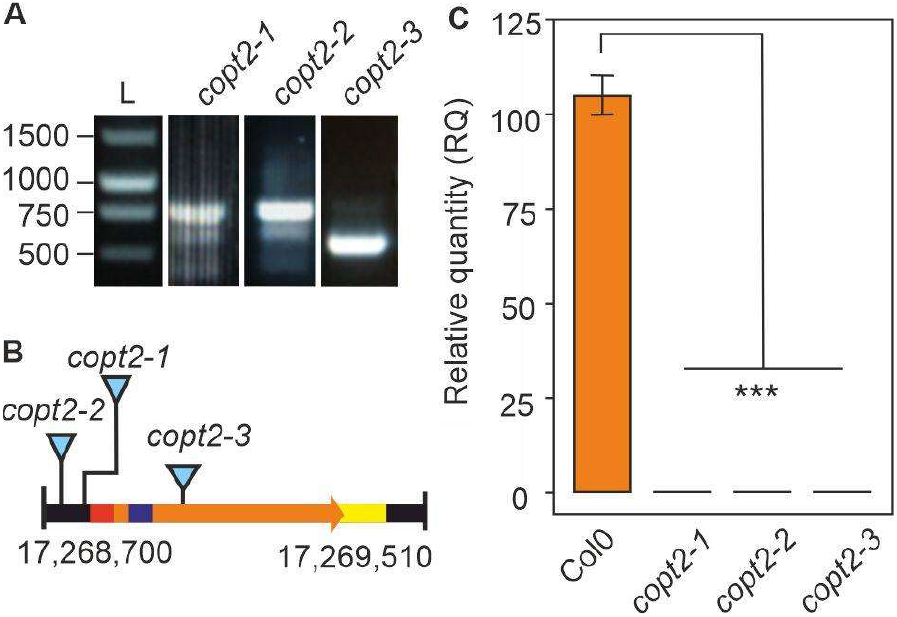
Characterisation of Arabidopsis *copt2* lines. (A) PCR analysis on genomic DNA using primers spanning T-DNA insert sites for the *copt2* mutants. L, DNA ladder. (B) Schematic showing locations of the T-DNA in the *copt2* mutants in chromosome 3. The bar colours correspond to specific regions: orange, *COPT2* gene sequence; red, 5’UTR; purple, translation start site; yellow, exon stop site (at junction) and 3’UTR. (C) qPCR analysis on cDNA from 2-week-old Arabidopsis rosette leaves. Relative quantity shows expression of *COPT2* relative to *actin*. Bars represent the mean of three plant replicates, *** denotes p-value > 0.001 following one-way ANOVA analysis.

### Root phenotypic response of Arabidopsis copt2 lines in response to Pd

In Arabidopsis, Cu availability strongly influences root architecture, with Cu excess leading to shortened primary roots, reduced branching, and disorganisation of the root meristem (Lequeux et al., 2010). Under control conditions (Fig. 3), two-week-old *copt2* plants exhibited significantly increased root length, fewer lateral branches, and fewer branches per mm. when compared to wild type plants. On agar plates containing 20 mM Pd, the root length of all three *copt2* lines was significantly reduced compared to the control condition, whereas wild type root length was unaffected. However, the root branching of the wild type was significantly decreased in the presence of Pd, while root branching of the *copt2* lines was less affected. These findings indicate that upon exposure to palladium, wild type roots exhibit stress-induced branching suppression from COPT2-mediated metal influx, whereas *copt2* mutants experience reduced primary root elongation due to pre-existing homeostatic stress, yet retain lateral branching because decreased intracellular Pd uptake protects their developing root primordia.

**Figure 3.**
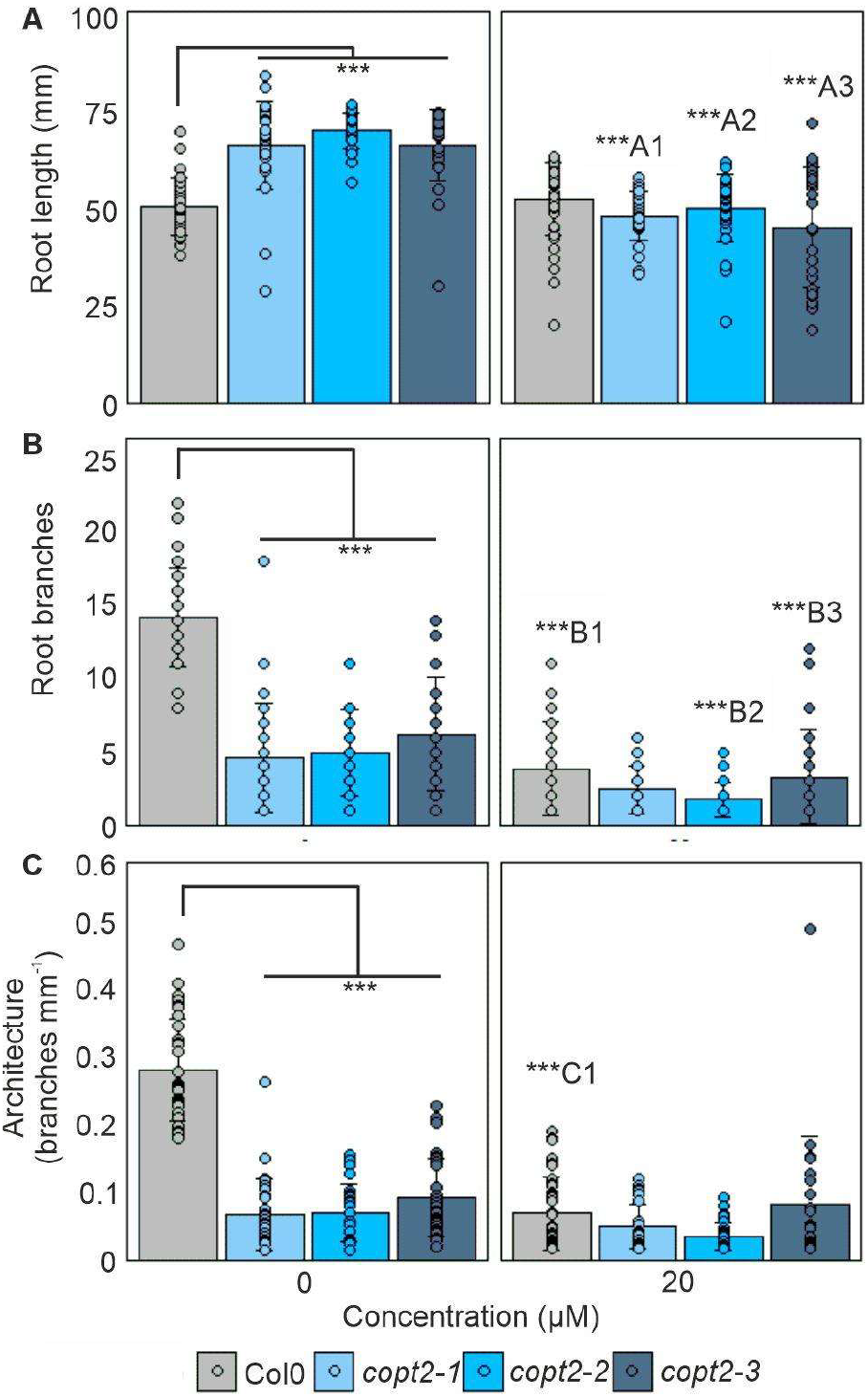
Measurement of Arabidopsis root architecture in presence of Pd. Root length, (A) Root branches, (B) and Architecture (C) in two-week-old plants grown on agar plates containing 1/2 MS media with and without 20 μM Pd. Data were analysed using a two-way ANOVA, with a p-value set to 0.02 due to deviations from assumptions in the data. Circular dots represent the distribution of raw data, while ‘whiskers’ indicate the upper and lower quartiles. Significant differences are denoted on the plots with asterisks (***: p < 0.001). Significance within each genotype between the 0 and 20 μM Pd conditions is identified by letters.

**Figure 4.**
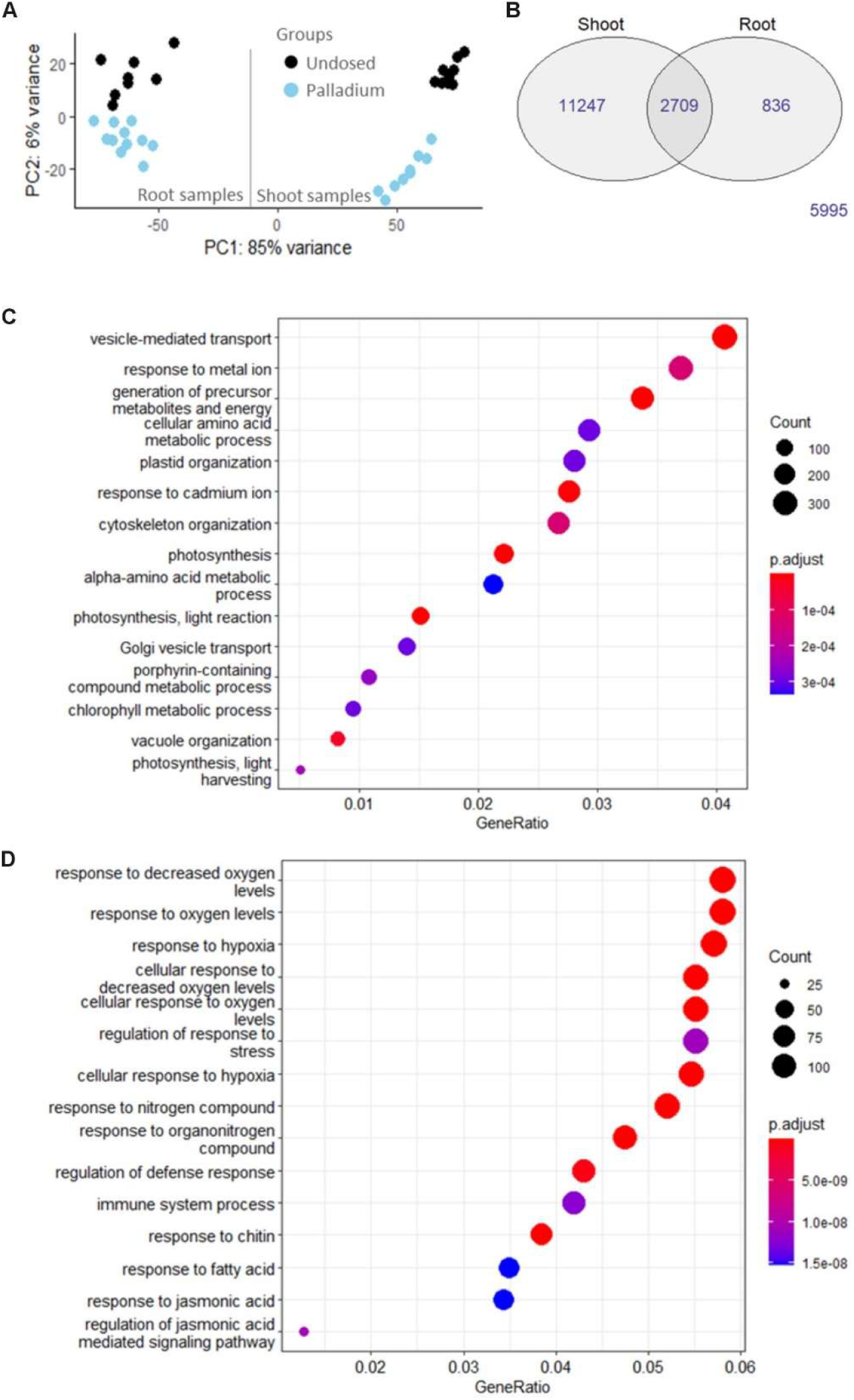
Analysis of transcriptomic data produced from hydroponically grown and Pd-dosed Arabidopsis plants. (A) Principal component analysis, (B) Number of differentially expressed genes (DEGs) present in shoot or root tissue, common to both tissue types, and those that were not differentially expressed. Dot plots illustrating the results of a gene ontology (GO) enrichment analysis. (C) Differentially expressed genes (DEGs) in shoot and (D) root tissue between undosed and Pd treated samples (log 2-fold change > 2; adjp < 0.05).

### Element concentration in Arabidopsis copt2 lines

Concentrations of Pd, Cu and iron (Fe) were measured in roots and shoots of five-week-old Arabidopsis plants dosed with 1 mM Pd (Fig. 5). A higher concentration than the 20 mM Pd in agar plates, was chosen here because we wanted to elicit a response in a shorter time frame (24h v two weeks) to avoid having developmental differences between treated and non-treated plants. Future studies could further optimise treatment duration and concentration. After 24h treatment with 1 mM Pd, a distinct brown colouration of the root tissues was observed (Fig. 5B), which remained despite multiple wash steps. Shoot tissues of the three *copt2* lines contained less Pd compared to wild type plants, significantly so for *copt2-2* and *copt2-3*. However, there was no difference in Pd concentration between wild type and *copt2* roots, and it is likely that adsorption of Pd to the outside of the root tissue masked analysis of root Pd-concentrations. Given the documented role of COPT2 in copper uptake (Mir, Pichtel and Hayat 2021, Sancenón et al. 2003; Loskarn et al., 2024), it was unexpected that Cu concentrations were unaltered in the *copt2* lines compared to the wild type. Interestingly, the *copt2* shoots contained less Fe, indicating possible crosstalk between Pd and Fe uptake pathways.

**Figure 5.**
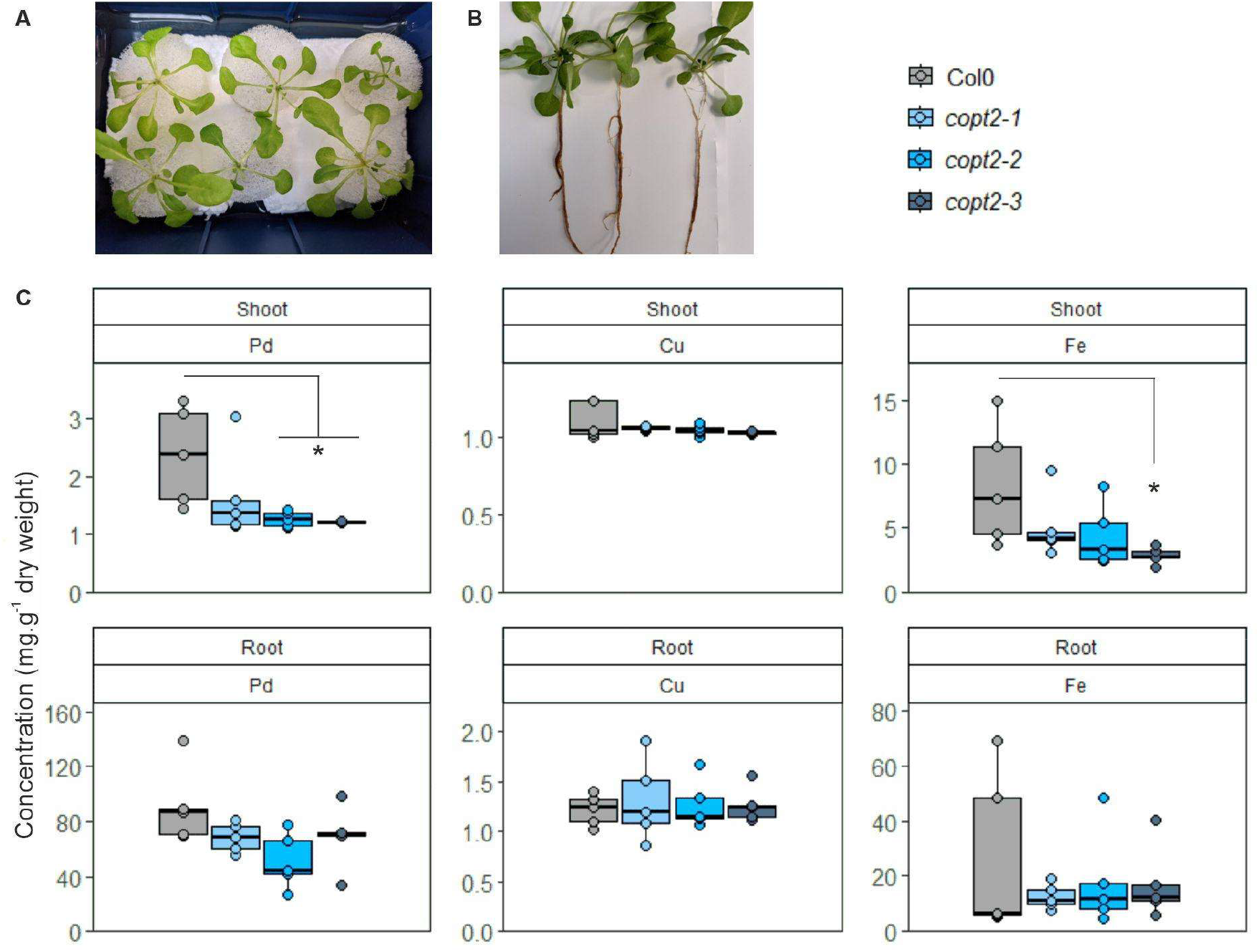
Element concentration in hydroponically grown Arabidopsis dosed with Pd. (A) Appearance of hydroponic system, and five-week-old Arabidopsis plants (B) immediately before treatment. Shoot (C) and root (D) metal contents 24h post treatment with 1 mM Pd. Element concentrations were measured using ICP_OES. Circular dots represent the distribution of raw data, while ‘whiskers’ indicate the upper and lower quartiles. Results were analysed using a two-way ANOVA. Significant differences between wild type Col0 and the three *copt2* lines are shown (* denotes p < 0.02; **, p < 0.01; ***, p < 0.001).

### Higher uptake of Pd results in an increase in oxidative stress in Col0

Metal stress can trigger the generation of free radicals and ROS (Prasad *et al*. 1999), and Tiwari *et al*. (2017) investigated the production of ROS in response to Au by staining with H_2_DCF in the root tip. In the presence of peroxides H_2_DCF is oxidised to the fluorescent product, dichlorofluorescein (DCF). To test if Pd elicited an oxidative response, plants were treated with 20 μM Pd for 24h (Fig. 6A), then stained with H_2_DCF, and fluorescence visualised. Figure 6B and C show that there was significantly less fluorescence produced in root tips of the *copt2-1* plants compared to the wild type controls.

**Figure 6.**
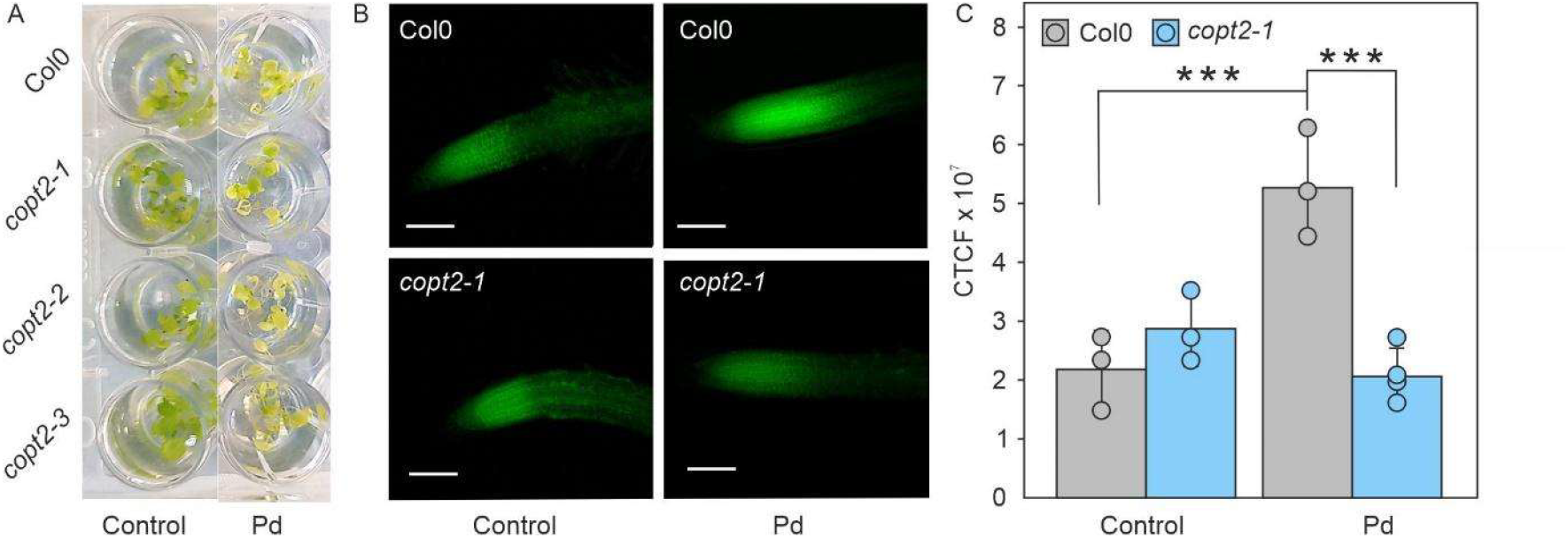
Pd-induced ROS production in Arabidopsis root tips. (A) Appearance of Arabidopsis before (left hand wells) and 24 hours after treatment (right hand wells) with 20 μM Pd. (B) Microscopic images showing ROS-linked fluorescence in Arabidopsis root tips. (C) Corrected total cell fluorescence (CTCF) in Arabidopsis root tips. Results are mean of three or more replicates, circular dots depict the raw data distribution, error bars indicate standard deviation. Results were analysed using a two-way ANOVA. Significant differences are shown (** denotes p < 0.01; ***, p < 0.001).

### Arabidopsis transcriptome response to Pd

To better understand the underlying strategies employed by plants to cope with Pd stress, we conducted a transcriptomics study on hydroponically grown Arabidopsis plants dosed with 1 mM Pd for 24 h. Principal component analysis (Fig. 4A) demonstrated distinct sample clustering within groups, with 85% of the variation ascribed to differences between tissue type (shoot vs. root) and 6% to the treatment (untreated vs. Pd-dosed). A Venn diagram illustrates the overlap of differentially expressed genes (DEGs) found across root and shoot tissues (Fig. 4B). Among these, 2,709 DEGs were identified in response to Pd across both tissue types, while 5,995 genes showed no significant differential expression. The top 20 most significantly up- or downregulated DEGs in shoot and root tissue in response to Pd treatment (log_2_ fold change ≥ 1 or ≤ −1, *p* < 0.05) are shown in Table S2.

Metal transporters such as COPTs, HMAs, and ZIPs (ZRT/IRT-like Proteins), alongside metal ligands such as metallothioneins, play crucial roles in metal uptake, homeostasis, and sequestration (Huang et al., 2024). Filtering the dataset for transporter activity revealed 26 DEGs in response to Pd (log_2_ fold change ≥ 1 or ≤ −1, *p* < 0.05) (Table S3). Although excluded from this table because it fell below the 2-fold cutoff, *COPT2* was significantly downregulated in shoot tissue in response to Pd (log_2_ fold change in shoots = −0.787) but showed no significant change in root tissue.

Because plant metal transporters display significant substrate promiscuity, it is likely that additional routes beyond COPT2 exist for Pd uptake. Arabidopsis Heavy Metal ATPases (HMAs) typically transport Zn, Cd, and/or Cu. In this study, *HMA7*, which facilitates Cu trafficking from the cytosol into Golgi apparatus storage reserves, was upregulated (shoot log_2_ fold change = 1.047, *p* < 0.001). A similar response was previously reported for *HMA7* following Cu treatment in bladder campion (*Silene vulgaris*) (Baloun et al., 2014). Conversely, *HMA2*, which encodes a Zn/Cd efflux transporter involved in root-to-shoot translocation, was downregulated in both roots and shoots under Pd exposure (log_2_ fold change = −2.924 and −1.637; *p* < 0.01 and *p* < 0.001, respectively), suggesting reduced Pd mobility and decreased accumulation in shoots.

In agreement with previous studies showing the downregulation of aquaporins in Arabidopsis plants treated with gold (Au) (Taylor et al., 2014), the aquaporin/arsenite transmembrane transporter 1 (*NLM1*, a NOD26-like major intrinsic protein) was significantly downregulated (log_2_ fold change = −2.036, *p* < 0.05). Gold is reported to covalently bind to cysteine residues and sulfhydryl groups of aquaporins (Niemietz and Tyerman, 2002; Preston et al., 2002), reducing root cell water permeability and overall water uptake. Given the chemical similarity between Au and Pd, Pd may target aquaporins through a similar mechanism.

Five out of the fifteen most enriched biological processes were related to oxygen stress (response to decreased oxygen levels, response to oxygen levels, response to hypoxia, cellular response to decreased oxygen levels, cellular response to oxygen levels, cellular response to hypoxia), plausibly an artifact of the hydroponic experimental set-up. While soil-based systems represent the ‘gold standard’ and are more representative of real-world scenarios, an advantage of hydroponics is that Pd remains in ionic form and is thus more readily bioavailable for studies focusing on *in planta* responses. In soil, Pd exists predominantly in unavailable forms, such as zero-valent metal. Furthermore, RNA extraction from soil-grown Arabidopsis roots is technically challenging, and the metal concentrations required for soil toxicity studies would be prohibitively expensive.

Dot plots were used to group the DEGs by broader biological pathways (Fig. S2). There was a significant enrichment of biological processes relating to general stress, defence, and immune responses (regulation of response to stress, regulation of defence response, immune system processes, response to jasmonic acid). At the gene level, glutathione transferases (GSTs) and glutamine synthetase (*GLN1;1*) were upregulated, reinforcing an enhanced oxidative stress defence and active detoxification response under Pd treatment.

Overall, the plant response to Pd mirrors established Zn/Cd and Cu detoxification pathways, indicating significant mechanistic overlap in uptake and sequestration systems. However, because Zn and Cd transport and detoxification have been studied far more extensively due to their agronomic relevance and toxicity, this existing bias likely shapes current interpretations of Pd uptake and tolerance mechanisms.

## Conclusions

Heterologous expression of *Arabidopsis* COPT2 in *S. cerevisiae* demonstrates its role in Pd uptake, as evidenced by increased Pd sensitivity in copper-regulation mutants. In Arabidopsis, characterisation of knock-out lines (*copt2-1, copt2-2, copt2-3*) further confirms COPT2-mediated Pd transport: loss of COPT2 leads to altered root architecture sensitivity under Pd exposure, significantly decreased Pd shoot accumulation, and reduced ROS generation in root tips under Pd stress.

Transcriptomic profiling reveals that Arabidopsis responds to Pd by downregulating root-to-shoot translocation transporters (*HMA2*) and aquaporins (*NLM1*), while upregulating cytosolic-to-Golgi sequestration routes (*HMA7*) alongside glutathione transferases and *GLN1;1*. Together, these findings indicate that COPT2 acts as a key influx pathway for Pd, and its loss reduces long-distance Pd transport to shoots, mitigating systemic oxidative stress through mechanisms that closely parallel established Cu and Zn/Cd homeostasis networks.

### Towards developing plants to recover PMGs from dilute mine wastes

Microorganisms are already employed in commercial processes to recover Au from relatively concentrated waste sources like waste electrical and electronic equipment (WEEE) recycling (Kwok, 2019). However, when it comes to mining dilute solutions of these valuable metals from larger areas such as mine tailings and road sweepings, plants offer additional benefits. These areas often suffer from soil degradation and pollution, resulting in low biodiversity and poor soil structure. Plants offer the potential to remediate soils and restore local ecology (Rylott and Bruce, 2022). Leveraging the resulting biomass for catalysis could enhance commercial value and render phytomining economically viable. To realise these advantages, it is essential for plants to efficiently take up and accumulate economically valuable levels of the target metal. Plant hyperaccumulators can concentrate specific metals, such as nickel, Cu, and Zn, in their tissues to levels many times higher than those found in the surrounding soil (Reeves *et al*., 2018). However, natural hyperaccumulators of gold and platinum group metals (PGMs) are lacking (Nemutandani *et al*., 2006).

Commercial application of phytomining for Au and PGMs also encounters significant challenges arising from the limited solubility and low uptake of these metals, as highlighted by Taylor *et al*. (2011) and Harumain *et al*. (2017a). To overcome these hurdles, the advancement of genetic engineering (GM) and plant synthetic biology techniques presents promising solutions to fine-tune the expression of genes responsible for metal sequestration, chelation, or reduction properties. This study contributes towards our understanding of the genetic basis behind the uptake, detoxification and accumulation of these elements in plants that is needed for this approach to succeed.

## Supplementary data

The following supplementary data are available at JXB online:

**Table S1**. PCR primers used in this study.

**Table S2**. Top 20 most significantly up or down regulated DEGs in shoot or root tissue in response to Pd treatment (log2fold change + > 1, p-value > 0.05).

**Table S3**. Significant DEGs encoding proteins with transporter activity in root tissue in response to Pd treatment.

## Acknowledgement

We acknowledge support of the Biotechnology and Biological Sciences Research Council: BB/X011232/1, BB/Y008456/1 and BB/M011151/1). And support from Ms Meg Stark, and colleagues at the University of York Bioscience Technology Facility.

